# Regulatory architecture controlling terminal differentiation of an interoceptive paraneuron in *C. elegans*

**DOI:** 10.64898/2026.05.28.728471

**Authors:** Hongzhu Ji, Berta Vidal, Elisabeth Conklin, Tergel Enkhtuvshin, Nathan E. Schroeder, Oliver Hobert

## Abstract

Interoceptive paraneurons are neuron-like cells located within internal epithelial cell surfaces that sense internal stimuli to evoke specific behavioral or physiological responses. The elucidation of terminal differentiation programs of paraneurons is expected to provide insights into how epithelial cells acquire neuron-like feature during development and possibly also over evolutionary time. We define here transcriptional programs that control the terminal differentiation of an interoceptive paraneuron class in the nematode *C. elegans*, called uv1. The uv1 cells sense mechanosensory inputs in the uterus and signal via the HSN neurons to modulate egg-laying behavior. We show that like in canonical neurons, the neuron-like secretory features of uv1 are controlled by a combination of CUT homeobox genes, while the combinatorial terminal gene battery that defines the unique functional features of uv1 is jointly controlled by a combination of at least three transcription factors, a LIM homeodomain (LIN-11), a SoxD (EGL-13) and a Pax family (EGL-38) protein. These factors act in a terminal selector-type manner to jointly co-regulate the many distinct uv1-paraneuron specific molecular features, such as sensory receptors, neuromodulatory receptors and neuropeptides, as well as uv1’s tyraminergic identity. Our findings demonstrate notable similarities in the dichotomous architecture of gene regulatory programs of neurons and paraneurons.

## INTRODUCTION

Paraneurons were defined by Fujita in the 1970s as cells that do not display an obvious neuronal morphology but that act in a neuron-like manner, being depolarized by external or internal signals and relaying these signals to the nervous system to control specific behavioral and physiological responses (Fujita, 1977; Fujita et al., 1988; Hobert, 2025). One prominent type of paraneurons are interoceptors located within internal epithelial surfaces, such as airways, the circulatory system or the intestine that sense specific internal cues and relay them to the nervous system. The functional similarities of neurons and paraneurons raises intriguing questions about what the definition of a neuron is, how cells acquire neuronal features during development, and also how neurons came into being in an evolutionary context, with paraneurons perhaps being illustrative of an intermediary step in the evolutionary trajectory of neurons (Hobert, 2025). One way to explore these issues is to (a) more precisely define the molecular composition of paraneurons and (b) to ask whether the gene regulatory architecture that controls the paraneuronal expression of such molecular features share any conceptually similarity with those of neurons.

The nematode *C. elegans* is particularly well suited to address these types of questions. First, cells that can be classified as paraneurons exist in *C. elegans* (Hobert, 2025). Second, the differentiation programs of neurons are particularly well-defined throughout the *C. elegans* nervous system, hence lending themselves to a deep comparison with paraneuronal differentiation programs that remain to be elucidated. Through the use of classic genetic loss of function analysis, several lessons have been learned about terminal differentiation programs in neuronal cell types of *C. elegans*: First, rather than being regulated in a piece-meal manner, large cohorts of terminal identity markers of a neuron are co-regulated by cooperating sets of transcription factor(s), called terminal selectors [reviewed in (Hobert, 2016)]. Terminal selectors do not act in isolation but cooperate in neuron type specific combinations, often acting in obligatory heteromeric complexes [reviewed in (Hobert, 2016)]. Second, many terminal selectors are homeodomain proteins, a possible reflection of their diverse protein-protein interaction surfaces (Hobert, 2021; Reilly et al., 2020). Third, while individual neuron type-specific gene batteries are controlled by terminal selectors, the expression of panneuronal identity features, shared by all neurons (such as synaptic vesicle proteins or neuropeptide processing machinery) are regulated by panneuronally expressed CUT homeodomain proteins that act in parallel to neuron-type specific terminal selector combinations (Leyva-Diaz and Hobert, 2022; Stefanakis et al., 2015). Hence, neuronal differentiation programs are characterized by a dichotomous gene regulatory architecture, in which two distinct gene regulatory routines act in parallel to control generic (panneuronal) and neuron type-specific features. We explore here whether the terminal differentiation programs of a paraneuron share a similar gene regulatory architecture.

The cells that fit most clearly the paraneuron definition in *C. elegans* are the four uv1 cells which derive from an epithelial cell layer in the uterus of the worm (Newman et al., 1996)(**Fig.1A**). uv1 cells are interoceptors that deploy TRP channels to sense internal pressure in the uterus, triggered by the presence of eggs in the uterus which stretches these epithelial cells (Collins et al., 2016; Jose et al., 2007; Yan et al., 2025)(**Fig.1A**). In response to mechanosensory signals, uv1 cells utilize a combination of neuropeptides, as well as the monoamine tyramine, directly synthesized by uv1, to modulate the activity of the HSN neurons, which directly innervate vulval musculature (Banerjee et al., 2017; Collins et al., 2016; Jose et al., 2007; Yan et al., 2025)(**Fig.1A**). uv1 activity is likely modulated by other signals as well, as inferred by the expression of monoaminergic, GABAergic and cholinergic metabotropic receptors (Fernandez et al., 2020) as well as other orphan GPCRs (Vidal et al., 2018).

**Figure 1.**
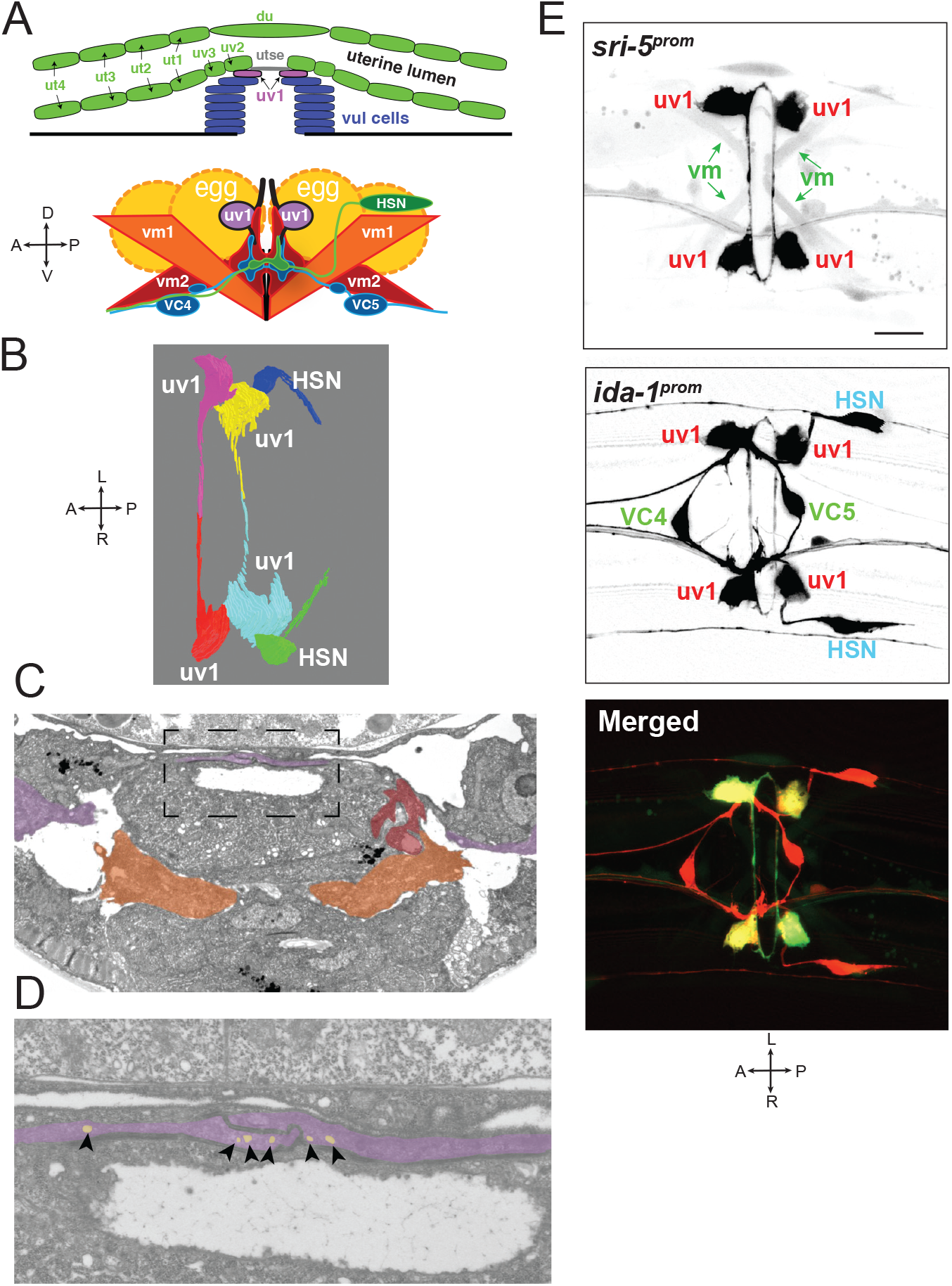
uv1 displays neuron-like features. **A:** Overview of structural elements of the uterus pertinent to the uv1 cells. Adapted from Collins et al., 2016 and Newman et al. 1996. **B:** Oblique dorsal 3D reconstruction view of vulval cells from EM data set. Also see supplemental video S1 and S2. **C,D:** EM from vulval region of late L4 *C. elegans* with pseudo-color overlay highlighting the uv1 (violet), the vulval muscles 1(orange) and vulval muscles 2 (red). Also see supplemental video S1 and S2. **D:** Inset from (C) showing junction of contralateral uv1 cells. Vesicles (arrowheads & pseudo-colored) are seen at various locations along the length of uv1 processes and some also at the soma (not shown). At least one vesicle is seen in 27/90 section out of which we reconstructed uv1. Also see supplemental video S1 and S2. **E:** Fluorescent reconstruction of *sri-5*^*prom*^*::gfp(otEx6406)* + *ida-1*^*prom*^*::rfp(vsIs269)* transgenic reporter strain. A: Anterior, P: Posterior, D: Dorsal, V: Ventral, L: Left, R: Right. Scale bar 10µm.

Previous work has provided substantial insights into the earlier patterning events that generate uterine and vulval cell types. The four uv1 cells are, together with their sister cells, the utse uterine epithelial cells, daughters of the π cell precursors (Gupta et al., 2012; Newman et al., 1996). uv1 fate is induced in a subset of the π cell descendants via an EGF signal emanating from neighboring vulval epithelial cells (VulF)(Chang et al., 1999; Gupta et al., 2012; Newman et al., 1996). Via an EGF-receptor and a *let-60/Ras*-dependent signaling cascade, this EGF signal induces the expression of the Pax transcription factor *egl-38/Pax2/5/8* in the nascent uv1 cells (Chamberlin et al., 1997; Chang et al., 1999; Rajakumar and Chamberlin, 2007; Webb Chasser et al., 2019). *egl-38* is required for the induction of several terminal markers of uv1 identity (neuropeptides *nlp-2* and *nlp-7*, as well as *ida-1*) (Rajakumar and Chamberlin, 2007; Webb Chasser et al., 2019). To what extent *egl-38* affects uv1 differentiation is not presently known, nor is it known whether *egl-38* is only transiently required to induce uv1 or is required after initial differentiation to maintain stable uv1 identity.

Two other transcription factors, the *lin-11* LIM homeobox gene, as well as the SoxD-type *egl-13* gene are also expressed in uv1 cells (Hanna-Rose and Han, 1999; Newman et al., 1999). Neither of these transcription factors appears to be required for each other’s expression and they may each be independent targets of EGFR signaling in uv1 (Hanna-Rose and Han, 1999; Newman et al., 1999; Rajakumar and Chamberlin, 2007). Due to the absence of proper differentiation markers at the time, uv1 differentiation has not been studied in *lin-11* mutants. Similarly, while it is known *egl-13/SoxD* affects π precursor cell proliferation and the proper cell fusion of the π-derived utse cells (Cinar et al., 2003), the impact *of egl-13/SoxD* on the differentiation of π-derived uv1 cells is not known either.

In this paper, we deploy a panel of cell identity markers to characterize the neuronal features of the uv1 paraneurons in more detail. We reconstruct the morphology of these cells and show that, like in neuronal cell types, uv1’s panneuronal features are controlled by CUT homeobox genes. We show that the *egl-38, lin-11* and *egl-13* transcription factors act akin to terminal selectors of neuronal cell types in that they co-regulate the entire battery of uv1-specific neuron-like features. Hence, our analysis not only provide more insights into the differentiation program of a paraneuron but also points toward striking similarities between neuronal and paraneuronal differentiation programs.

## MATERIAL AND METHODS

### Strains

A list of strains used in this study is provided in Table S1.

### *C. elegans* genome-engineered strains

#### lin-11 null alleles

Due to issues with their mating efficiency, multiple molecularly identical *lin-11* null alleles, removing the entire coding region, were generated by injecting separate stains carrying different uv1 reporters. CRISPR/Cas9 genome engineering was performed as described in (Eroglu et al., 2023). The following crRNA and ssODN sequences were used:

crRNA1: ATTGAGAAGGGAGTAAAAGG crRNA2: CGTGGAATACTCCTGTATGT

ssODN: TTCGTGGTCGTTCTTCTTCTTCTTCTCCTCCTCCTTACAGGAGTATTCCACGTTCGTGTAGTTTTTCTT C

#### *lin-11(ot1834)* and *egl-38(ot1835)* AID alleles

*lin-11* and *egl-38* loci were tagged at the C-terminus with the sequence GSGGSGGTGGSG::mIAA7::wrmScarlet-I3::mIAA7 using CRISPR/Cas9 genome engineering as described in (Eroglu et al., 2023). The following crRNA sequences were used:

*lin-11(ot1834)* crRNA: agacttgggaaaaccaactc

*egl-38(ot1835)* crRNA: attctaccacaaaactattg

For the preparation of the ssODN, since PCR amplification of a fragment containing two identical mIAA7 sequences was problematic we ordered a synthetic gene fragment with two different synonymous mIAA7 sequences which could be amplified by PCR without problems.

#### *nas-22(ot1844) and exc-9(ot1866)* reporter alleles

*nas-22 and exc-9* loci were tagged at the C-terminus with the sequence SL2::GFP::H2B using CRISPR/Cas9 genome engineering as described in (Eroglu et al., 2023). The following crRNA sequences were used:

*nas-22(ot1844)* crRNA: gtcaaactccaacaattgtc

*exc-9(ot1866)* crRNA: tcaactgcttctccacgttc

#### *unc-31(syb6138)* and *unc-13(syb6325)* reporter alleles

*unc-31* and *unc-13* loci were tagged at the C-terminus with a SL2::GFP::H2B tag using CRISPR/Cas9 genome engineering. These strains were generated by SunyBiotech.

### Conditional LIN-11 and EGL-38 protein degradation

We used conditional protein depletion with a modified auxin-inducible degradation system, AID2 (Hills-Muckey et al., 2022; Sepers et al., 2022). mIAA7-tagged proteins are conditionally degraded when exposed to 5-phenyl-indole-3-acetic acid (5-Ph-IAA) in the presence of AtTIR1^F79G^. To generate the experimental strains, the conditional alleles *lin-11(ot1834[lin-11::mIAA7::Scarlet-I3::mIAA7]*) and *egl-38(ot1835[egl-38::mIAA7::Scarlet-I3::mIAA7])* were crossed with *osIs182[eft-3p::TIR1(F79G)]*, which expresses AtTIR1^F79G^ ubiquitously. Conditional LIN-11 and EGL-38 protein degradation was performed on NGM plates containing 100 µM 5-Ph-IAA that were prepared using a previously described protocol (Sural et al., 2024). Animals were transferred to 5-Ph-IAA plates at the mid L4 stage and imaged 1 day later. Control plates had the same volume of solvent alone (ethanol). All plates were stored in the dark for the entire duration of the experiment.

### Cell identification

Expression of *gfp-*based reporter genes in uv1 was confirmed with a *bona fide* uv1 marker, *ida-1::mCherry*, present in the background. In mutant backgrounds, a loss of *gfp-*based marker gene expression was usually accompanied by a loss of *ida-1::mCherry* as well, and in these cases we assessed loss of signal based on presumptive cell position of uv1 and other landmarks around the cell.

### Fluorescent microscopy

Worms were anesthetized using 100 mM sodium azide (NaN3) and mounted on 5% agarose pads on glass slides. z-stack images (0.5-1 μm thick) were acquired using a Fluar 40x/1.30 Oil objective on a Zeiss compound microscope (Imager Z2) with ZEN Blue software. Maximum intensity projections of 2-30 slices were generated with Fiji/ImageJ software (Schindelin et al., 2012). In many images the slices are focused around the uv1 cells, leaving other cells either out of focus or not visible at all and therefore giving the false impression that marker expression in other cells may be variable.

### Electron microscopy

Archived electron microscopy negatives and prints of a late L4 hermaphrodite, originally collected at the Medical Research Council (MRC) and previously published in (Newman et al., 1996) were digitized at high resolution by the Center for *C. elegans* Anatomy. Negatives were aligned using TrakEM2 (Cardona et al., 2012) and imported into VAST (Berger et al., 2018) for segmentation. Identification of individual cells was made by referencing original print annotations and corresponding MRC notebooks. 3D models .obj files were generated in VAST and imported into Blender for rendering following a remesh modifier and Laplacian smoothing.

### Statistical analysis

Data points for all biological replicates and sample sizes (N) are displayed in the figures. Statistical tests used in each figure and the corresponding P values are listed in the figure legends. P < 0.05 was considered to be statistically significant. All statistical tests and plotting were performed on GraphPad Prism 10.

## RESULTS

### Morphology of uv1 cells

The morphology and shape of uv1 cells have not been described in detail before. We reconstructed the morphology of the four uv1 cells using available electron micrographical images (**Fig.1B-D; Suppl. Video S1 & S2**), as well as optical reconstruction of confocal image stacks of a transgenic strain that expresses an uv1-restricted marker, *sri-5::gfp* (Vidal et al., 2018), and *ida-1::rfp*, a marker that is expressed in the entire egg-laying circuit, including uv1, VC4/5 and HSN (Banerjee et al., 2017; Zahn et al., 2001)(**FIG.1E**). Together, these different visualization approaches reveal a neuron-like morphology of the uv1 cells, extending thin processes along the lining of the uterine opening to the vulva. At this point it remains unclear whether it may be the stretch of these thin processes or pressure on the uv1 soma that can be made responsible for the mechanosensory function of uv1. The EM data also suggests the presence of vesicles along the length of uv1 processes (**Fig.1D**). However, the vesicles do not have the appearance of dense-core vesicles, and they are not clustered at specific release site. Taken together, the interoceptive function of uv1, its unusual morphology, but apparent ability to communicate with other neurons allow the uv1 cells to be classified as “paraneurons”, even though their previous naming as “neuroendocrine” remains equally valid.

### uv1 expresses panneuronal features in a CUT gene-dependent manner

Previous reporter gene studies, using fosmid-based reporter transgenes had already noted expression of several, panneuronally expressed synaptic vesicle and neuropeptide-processing/secretory proteins in the uv1 cells (Stefanakis et al., 2015). We corroborated this observation using both improved reagents (engineered reporter alleles which tag the endogenous gene locus vs fosmid-based transgenes) and additional markers (**Table 1**). This includes two CRISPR/Cas9-engineered reporter alleles that we generated specifically for this study, reporting on expression of *unc-13* and *unc-31* (see Methods). We found that markers of the synaptic vesicle cycle (*unc-18, unc-13, ric-4/SNAP25, rab-3*), vesicular transport (*unc-104/kinesin)* as well as genes involved in neuropeptide processing and signaling (*egl-3/PC2, unc-31/CAPS*) are expressed in uv1 (**Fig.2A**). We also found that the voltage gated calcium channel, *unc-2*, for which we generated a novel reporter allele (M. Boeglin, pers. comm.) is expressed in uv1. The NeuroPAL reporter transgene, which contains the so-called UPN panneuronal driver, a composite of *cis-*regulatory elements from several panneuronal genes (Yemini et al., 2021), is also expressed in uv1 (**Fig.2A**).

**Table 1.**
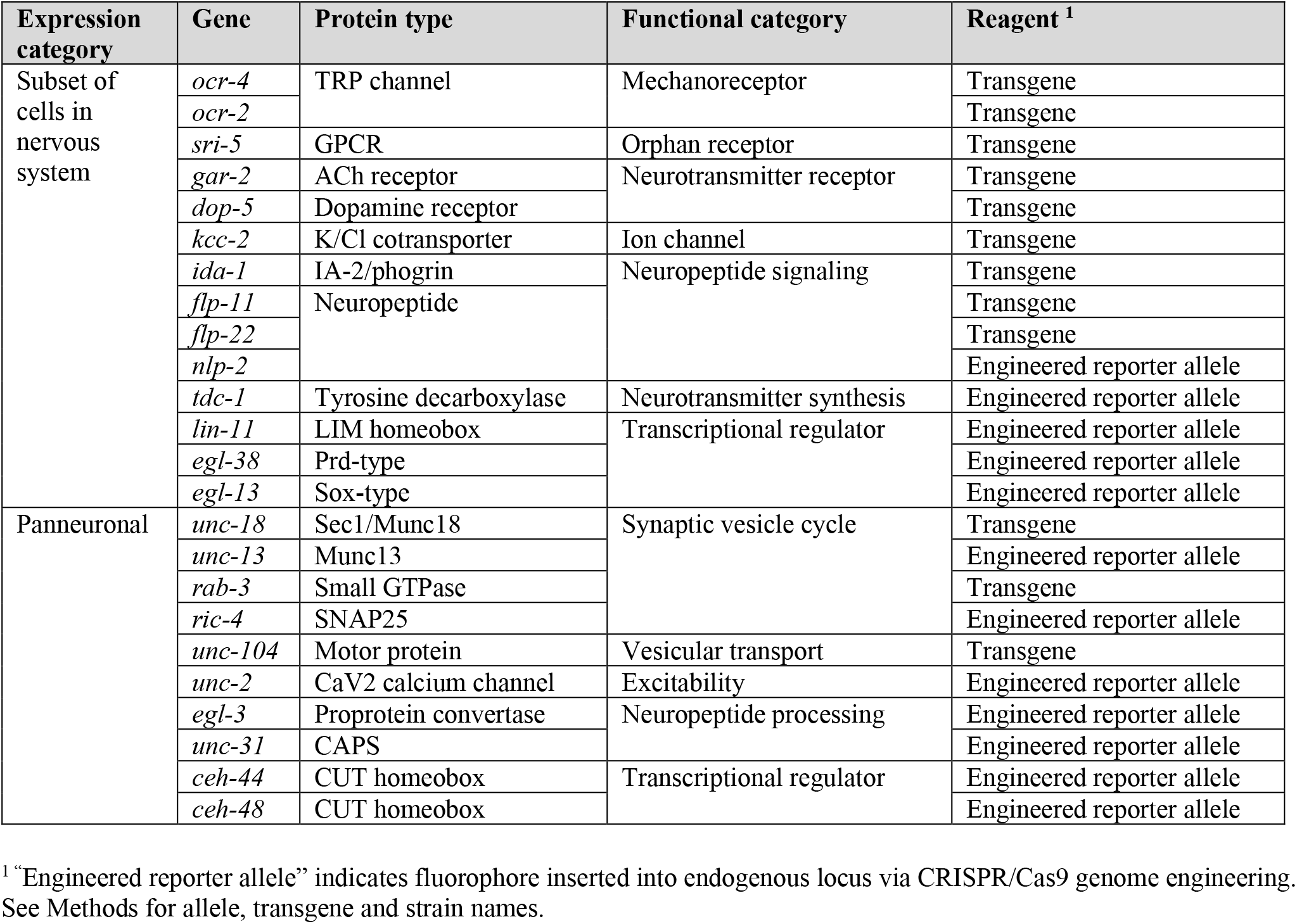
uv1 cell identity markers used in this study.

| Expression category | Gene | Protein type | Functional category | Reagent <sup>1</sup> |
| --- | --- | --- | --- | --- |
| Subset of cells in nervous system | *ocr-4* | TRP channel | Mechanoreceptor | Transgene |
|  | *ocr-2* |  |  | Transgene |
|  | *sri-5* | GPCR | Orphan receptor | Transgene |
|  | *gar-2* | ACh receptor | Neurotransmitter receptor | Transgene |
|  | *dop-5* | Dopamine receptor |  | Transgene |
|  | *kcc-2* | K/Cl cotransporter | Ion channel | Transgene |
|  | *ida-1* | IA-2/phogrin | Neuropeptide signaling | Transgene |
|  | *flp-11* | Neuropeptide |  | Transgene |
|  | *flp-22* |  |  | Transgene |
|  | *nlp-2* |  |  | Engineered reporter allele |
|  | *tdc-1* | Tyrosine decarboxylase | Neurotransmitter synthesis | Engineered reporter allele |
|  | *lin-11* | LIM homeobox | Transcriptional regulator | Engineered reporter allele |
|  | *egl-38* | Prd-type |  | Engineered reporter allele |
|  | *egl-13* | Sox-type |  | Engineered reporter allele |
| Panneuronal | *unc-18* | Sec1/Munc18 | Synaptic vesicle cycle | Transgene |
|  | *unc-13* | Munc13 |  | Engineered reporter allele |
|  | *rab-3* | Small GTPase |  | Transgene |
|  | *ric-4* | SNAP25 |  | Engineered reporter allele |
|  | *unc-104* | Motor protein | Vesicular transport | Transgene |
|  | *unc-2* | CaV2 calcium channel | Excitability | Engineered reporter allele |
|  | *egl-3* | Proprotein convertase | Neuropeptide processing | Engineered reporter allele |
|  | *unc-31* | CAPS |  | Engineered reporter allele |
|  | *ceh-44* | CUT homeobox | Transcriptional regulator | Engineered reporter allele |
|  | *ceh-48* | CUT homeobox |  | Engineered reporter allele |

<sup>1</sup> “Engineered reporter allele” indicates fluorophore inserted into endogenous locus via CRISPR/Cas9 genome engineering. See Methods for allele, transgene and strain names.

**Figure 2.**
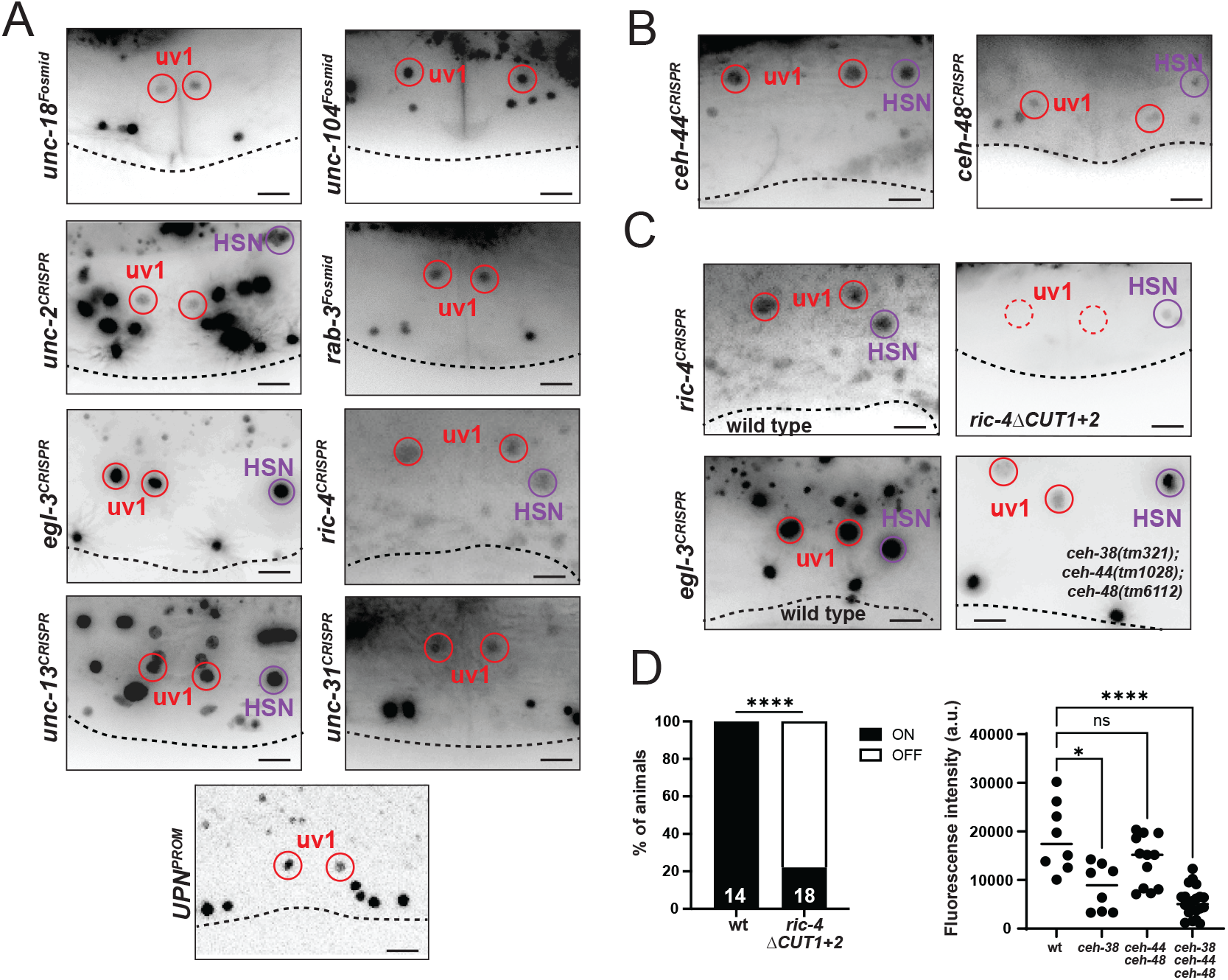
Expression and regulation of panneuronal markers in uv1cells. **A:** Expression of panneuronal markers in uv1cells. Reporter genes used are CRISPR/Cas9-engineered reporter alleles for *unc-2(ot1614[unc-2::SL2::gfp::h2b]), egl-3(syb4478[egl-3::SL2::gfp::h2b]), unc-13(syb6325[unc-13::SL2::gfp::h2b]), unc-31(syb6138[unc-31::SL2::gfp::h2b]), ric-4(syb2878[ric-4::SL2::gfp::h2b])* and transgenic reporters for *unc-18(otEx5925), unc-104(otEx4555), rab-3(otIs498)* and *UPN(otIs669)*. **B:** Expression of panneuronal CUT homeobox genes *ceh-44* and *ceh-48* in uv1 cells. Reporter genes used are CRISPR/Cas9-engineered translational *gfp* reporter fusions for *ceh-44(ot1015[ceh-44::gfp])* and *ceh-48(ot1126[ceh-48::gfp])*. **C:** *ric-4(syb2828)* reporter expression in CUT-site deleted allele (*ot1123 ot1179*) (top panel). *egl-3(syb4478)* reporter expression in *ceh-38(tm321), ceh-44(tm1028)* and *ceh-48(tm6112)* CUT homeodomain triple mutant (bottom panel). **D:** Quantification of *ric-4* reporter signal from panel C, statistical analysis was performed using Fisher’s exact test (left) and Kruskal-Wallis test (right), ****P < 0.0001. “ON” and “OFF” refers to reporter signal. Animals in all panels were imaged at the young adult stage and uv1 cells were identified by co-localization with a *tdc-1* or *ida-1* red marker in the background (not shown). All images are approximate lateral views with anterior to the left and ventral down. Dashed lines delineate ventral outline of worms. Scale bars 10µm.

Panneuronally expressed CUT homeobox genes act in neurons to control panneuronal gene expression (Leyva-Diaz and Hobert, 2022). Using CRISPR/Cas9-engineered reporter alleles, we observe expression of the two otherwise neuron-restricted CUT homeodomain proteins CEH-44 and CEH-48 in uv1, in addition to the ubiquitously expressed CEH-38 homeodomain protein (**Fig.2B**).

We demonstrated the requirement of these CUT homeobox genes for expression of panneuronal features in uv1 in two independent manners. First, we assessed expression of the panneuronal marker *egl-3* in CUT homeobox single, double and triple mutant background. As reported in the nervous system (Leyva-Diaz and Hobert, 2022), we observe a reduction of *egl-3* expression (**Fig.2C,D**). Second, we examined expression of a reporter gene-tagged *ric-4/SNAP25* locus in which the CUT homeodomain have been deleted through CRISPR/Cas9 genome engineering. This *cis*-regulatory allele shows a reduction of expression in the nervous system (Leyva-Diaz and Hobert, 2022) and we observe a reduction of expression in the uv1 cells as well (**Fig.2C,D**).

### The transcriptional regulatory signature of uv1

As a first step to understand uv1 cell differentiation, we set out to examine the expression of cell type-specifically expressed transcription factors in these cells. Previous transgenic reporter gene studies had shown the combinatorial expression of three transcription factors, the LIM homeodomain protein LIN-11, the Sox protein EGL-13 and Pax protein EGL-38 in uv1 (Hanna-Rose and Han, 1999; Newman et al., 1999; Rajakumar and Chamberlin, 2007). Since these observations were made with reporter transgenes that captured only part of the surrounding regions of each locus, we sought to validate these findings using fosmid-based reporters as well as CRISPR/Cas9-engineered reporter alleles. These reagents corroborated continuous co-expression of these three transcription factors in the 4 uv1 paraneurons throughout the life of these cells (**Fig.3**). Since their expression does not appear to overlap elsewhere in the animal, *lin-11, egl-38* and *egl-13* define a unique regulatory signature within the uv1 cells.

**Figure 3.**
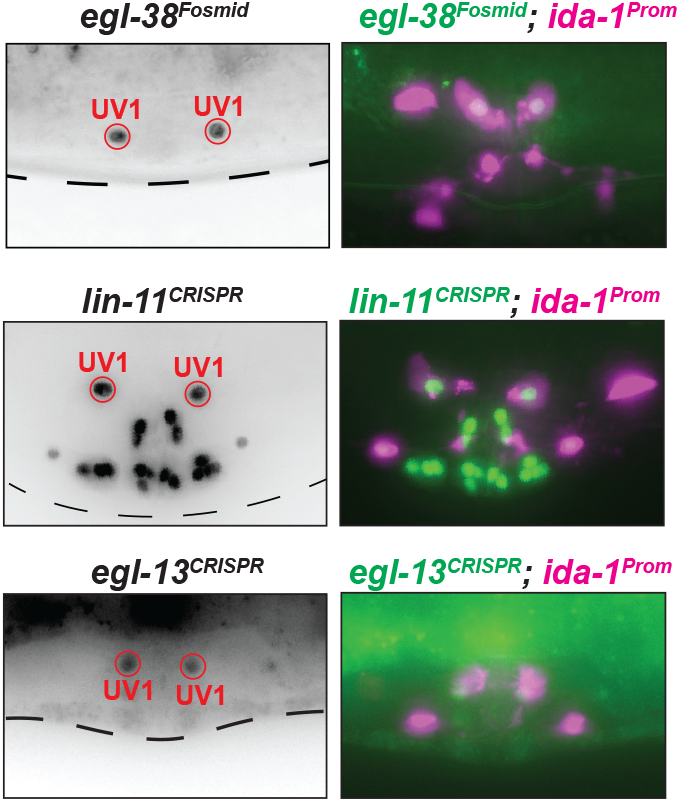
A combinatorial transcription factor code for uv1 Expression of cell-specific regulatory factors *egl-13, lin-11and egl-38* in uv1. All strains have *ida-1prom::rfp* reporter in the background to assess uv1 position. Reporter genes used are CRISPR/Cas9-engineered reporter alleles for *lin-11(ot958), egl-13(dev199)* and a transgenic fosmid reporter for *egl-38(wgIs171)*. A *mScarlet-*tagged *egl-38* locus (**Fig.5**) shows the same expression in uv1 as the fosmid-based reporter.

### *lin-11* coordinately regulates a large cohort of uv1 identity markers

We first tested the involvement of *lin-11* in uv1 differentiation. Previous work examining *lin-11* gene function had used either hypomorphic alleles or a presumed *lin-11* null allele that eliminated only parts of the locus and also eliminated two adjacent loci (Young et al., 2022). We generated unambiguous molecular null alleles of *lin-11* through the elimination of all coding region of the locus using the CRISPR/Cas9 system (Fernandez et al., 2025). To assess the effect of *lin-11* removal, we assessed the expression a large array of terminal identity markers of uv1 that have been described in the literature over the last few years (summarized in **Table 1**). These include a wide-range of effector genes that are, in addition to uv1, also expressed selectively in specific neuron classes and therefore attest to the neuron-like molecular identity of uv1. These markers include a mechanosensory TRP channels (*ocr-4*)(Jose et al., 2007; Upadhyay et al., 2016), three GPCRs (*gar-2, dop-5*)(Fernandez et al., 2020; Vidal et al., 2018), two of which metabotropic neurotransmitter receptors (ACh for *gar-2*, monoamine for *dop-5*), the ligand-gated ion channel *lgc-55* (Collins et al., 2016), four neuropeptides (*flp-11, flp-22, nlp-2, nlp-7*)(Banerjee et al., 2017; Kim and Li, 2004; Van der Auwera et al., 2020), a dense core vesicle component involved in neuropeptide release (*ida-1*) (Banerjee et al., 2017; Zahn et al., 2001), the tyramine biosynthetic enzyme *tdc-1* (Alkema et al., 2005) and a potassium chloride cotransporter, *kcc-2* (Tanis et al., 2009). We found that the expression of every single marker is essentially eradicated upon genetic elimination of *lin-11*, achieved through CRISPR/Cas9 genome engineering (**Fig.4A**).

**Figure 4.**
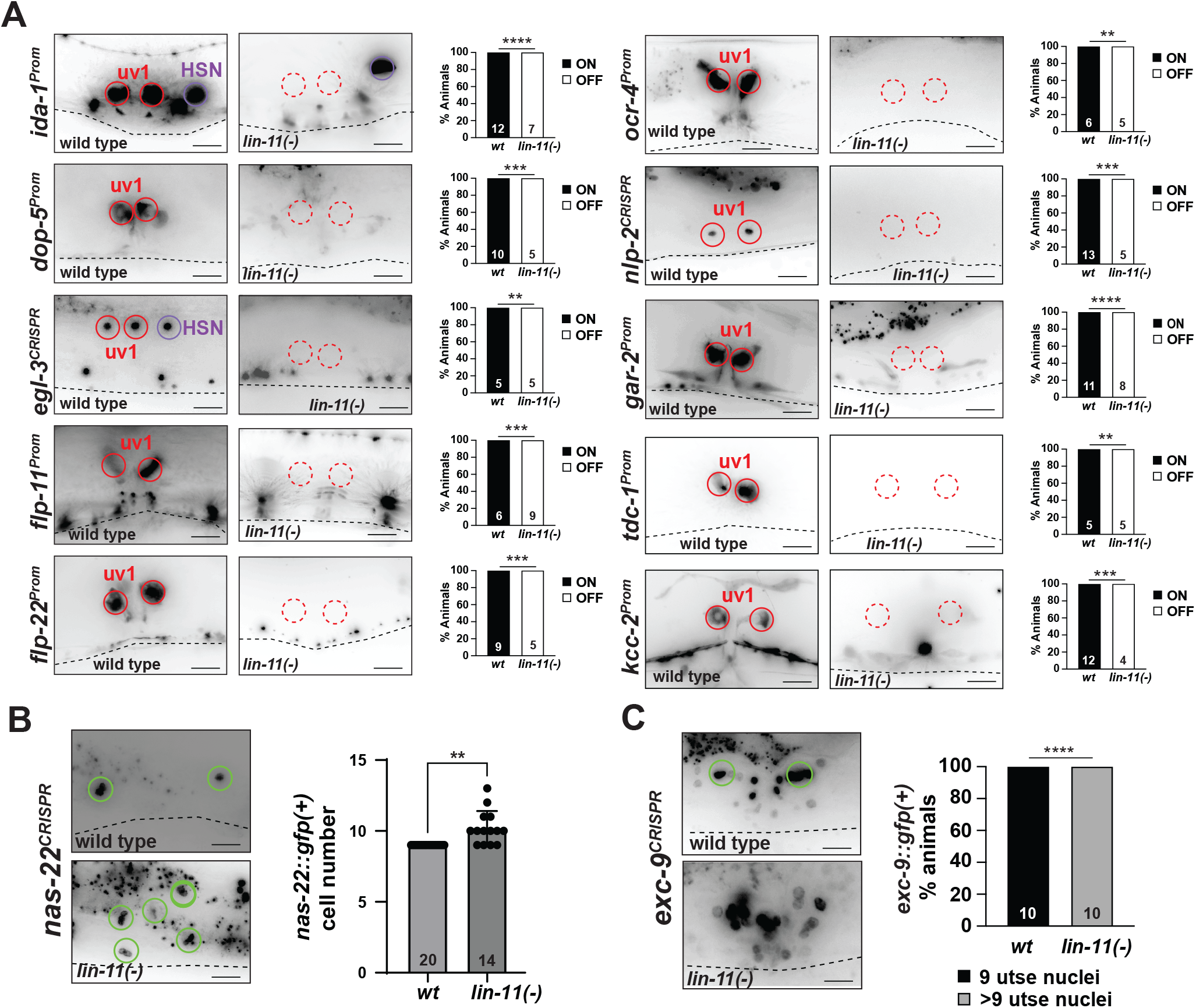
Loss of *lin-11* results in an uv1 to utse transformation. **A:** Effect of *lin-11* null mutant on uv1 markers. All panels show molecularly identical *lin-11* null alleles *(ot1867, ot1868, ot1869, ot1870, ot1871, ot1872, ot1873, ot1874, ot1875, ot1876, ot1877, ot1878, ot1845)* generated independently in different reporter backgrounds with identical crRNA and ssODN reagents. Reporter genes used are CRISPR/Cas9-engineered reporter alleles for *egl-3(syb4478), nlp-2(syb5697)* and transgenic reporters for *ida-1(vsIs269), dop-5(vsIs254), flp-11(ynIs40), flp-22(ynIs50), gar-2(vsEx912), ocr-4(kyEx581), tdc-1(zfIs10)* and *kcc-2(kvIs136)*. “ON” and “OFF” refers to reporter signal in wildtype versus *lin-11* mutant background. **B, C:** *lin-11* null mutants have additional utse cells as seen with *nas-22(ot1844[nas-22::SL2::gfp::h2b])* (panel B) and *exc-9(ot1866[exc-9::SL2::gfp::h2b])* CRISPR/Cas9-engineered reporter alleles (panel C). Animals in all panels were imaged at the young adult stage. All images are approximate lateral views with anterior to the left and ventral down. Dashed lines delineate ventral outline of worms. Scale bars 10µm. Statistical analysis was performed using Fisher’s exact test (A, C) and Welch’s t-test (B), **P < 0.01, ***P < 0.001, ****P < 0.0001.

To assess the possibility that in *lin-11* mutants, the uv1 paraneurons differentiate instead into the fate of their sister cells, the uterine epithelial utse cells, we developed cellular fate markers for this cell type. Specially, we selected the astacin-like protease NAS-22 and the LIM-only domain protein EXC-9 as candidates, based on pervious transgenic reporter gene studies (McKay et al., 2003; Tong and Buechner, 2008). We inserted an SL2::GFP::H2B tag at the 3’end of the respective coding regions in the genome and indeed observed restricted, nuclear localized expression in the utse cells and no expression in uv1 (**Fig.4B,C**). In *lin-11* null mutants, we observed additional cells expressing both of these markers, consistent with the possibility of an uv1 to utse cell fate change (**Fig.4B,C**).

### Temporally controlled *lin-11* removal demonstrates protein function in mature uv1 cells

Since *lin-11* is not only expressed during and after uv1 differentiation, but already in their precursor cells, the πcells, as well as in the vulF cells required to induce uv1 fate (Newman et al., 1999), we tested whether we can tie *lin-11* function more selectively to the induction of the terminally differentiation state of uv1. To this end we deployed the auxin-dependent degron system. The mScarlet-I3 fluorophore with which we had tagged the *lin-11* locus contains two separate mIAA7 degrons at each end of the mScarlet-I3 sequence. Using a ubiquitously expressed TIR1^F79G^-driver line, we exposed *lin-11::mIAA7::mScarlet-I3::mIAA7* animals at the mid-L4 stage to auxin. Taking into account the time it takes for auxin to induce LIN-11 degradation and the time it takes to then observe an effect on LIN-11 downstream target expression, we infer that this timing of removal probes LIN-11 requirement for proper uv1 differentiation. Using four different terminal uv1 markers (*flp-*22, *unc-2, tdc-1* and *unc-13)*, we found that LIN-11 removal affects expression of all these markers (**Fig.5A**). We conclude that LIN-11 is required for the initiation/maintenance of the terminal uv1 differentiation program.

**Figure 5.**
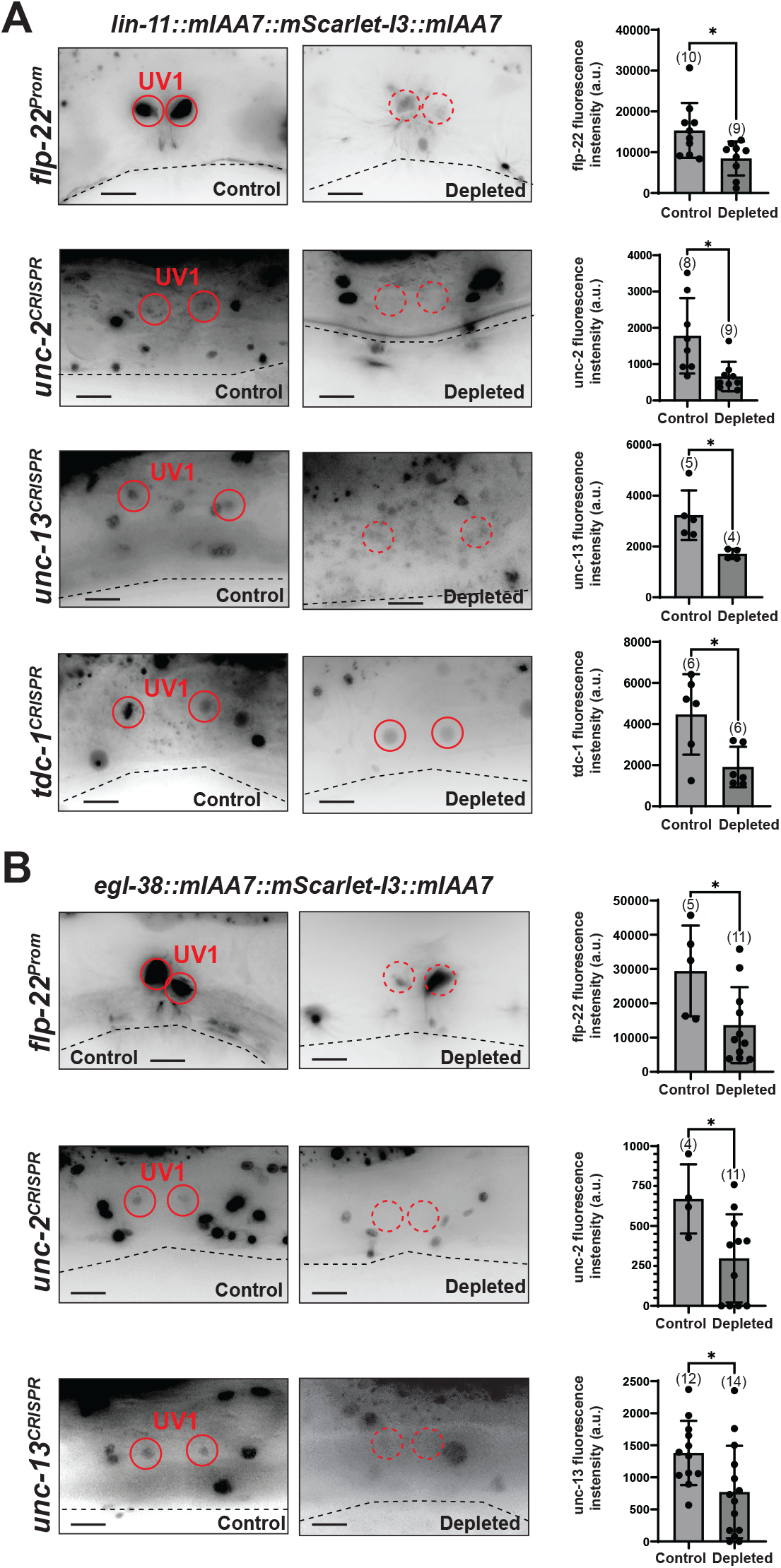
Temporally controlled LIN-11 & EGL-38 removal disrupts uv1 differentiation. *lin-11(ot1834[lin-11::mIAA7::mScarlet-I3::mIAA])* (panel A) and *egl-38(ot1835[egl-38::mIAA7::mScarlet-I3::mIAA])* (panel B) were temporally depleted with the AIDv2 system (Hills-Muckey et al., 2022). Worms at the mid L4 stage were transferred onto ethanol-(Control) or 5-Ph-IAA-coated plates (Depleted) and scored 1 day later. Worms were expressing TIR1^F79G^ under the ubiquitous *eft-3* promoter (*osIs182*). Reporter genes used are CRISPR/Cas9-engineered reporter alleles for *unc-2(ot1614), unc-13(syb6325), tdc-1(syb7768)* and transgenic reporter for *flp-22(ynIs50)*. All images are approximate lateral views with anterior to the left and ventral down. Maximum intensity projections of limited image slices are shown, which makes expression in other cells seem more variable than it really is; oftentimes the other cells are not within the image slices used. Dashed lines delineate ventral outline of worms. Scale bars 10µm. Statistical analysis was performed using Welch’s t-test, *P < 0.05.

### The Paired-box EGL-38 protein affects uv1 differentiation

Using three uv1 markers (*nlp-2, nlp-7* and *ida-1*), an effect of a hypomorphic allele of the Pax2/5/8-ortholog *egl-38* on uv1 differentiation has been reported in earlier reports (Rajakumar and Chamberlin, 2007; Webb Chasser et al., 2019). An assessment of *egl-38* function in uv1 is, however, complicated by the fact that *egl-38* is known to be already required in the vulF cells to produce the EGF signal that is essential for uv1 fate specification (Chamberlin et al., 1997; Chang et al., 1999). Constitutive activation of this signal has been used to mask this earlier function of *egl-38* and focus on the function of *egl-38* in uv1 (Rajakumar and Chamberlin, 2007), but it is difficult to exclude the possibility that the uv1 differentiation defects observed in this genetic background are entirely independent of earlier *egl-38* function in vulF. To circumvent this earlier function of *egl-38*, we used an alternative strategy, again making use of the auxin-inducible degron system. A strain in which mIAA7 degrons were inserted, together with mScarletI3, at the 3’end of the *egl-38* locus was subjected to auxin treatment at the L4 stage. We found that such removal affects the expression of three terminal uv1 markers, *flp-22, unc-2* and *unc-13* (**Fig.5B**). As with LIN-11, we interpret these results to mean that EGL-38 acts during the initiation/maintenance phase of the terminal uv1 differentiation program.

### The SoxD transcription factor *egl-13* also affects uv1 differentiation

*egl-13/SoxD* had previously been shown to be expressed in uv1 (Hanna-Rose and Han, 1999) but its function in uv1 differentiation has not previously been examined. To assess a potential role of *egl-13*, we utilized the previously described *ku194* null allele of *egl-13* (Hanna-Rose and Han, 1999) and crossed nine of the uv1 cell identity markers described above into this mutant background. We found that expression of each of these markers is lost in *egl-13* mutants (**Fig.6A**). Similarly, the panneuronal uv1 color of the NeuroPAL transgene is lost (**Fig.6A**).

**Figure 6.**
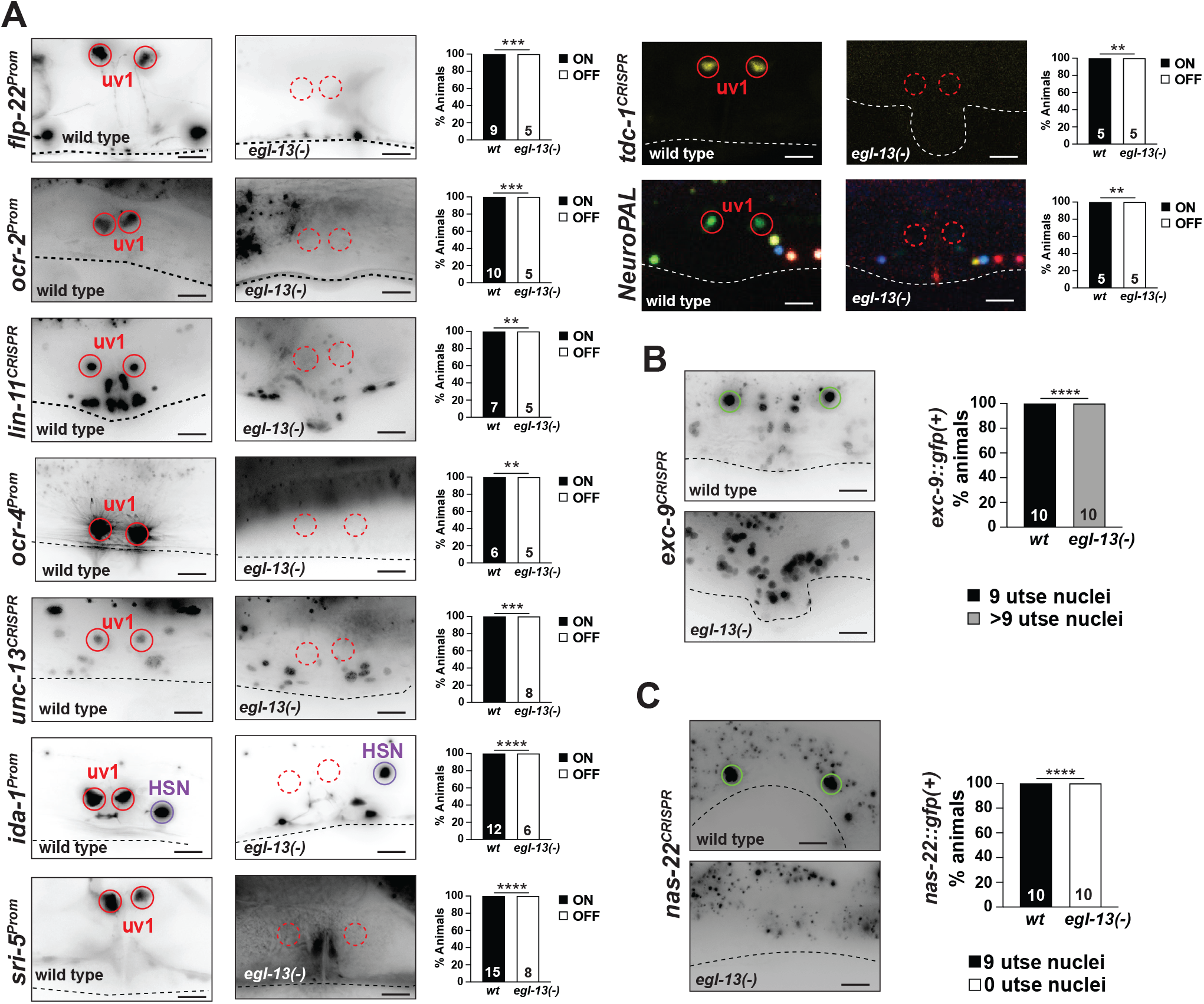
The SoxD transcription factor *egl-13* affects uv1 differentiation. **A:** Effect of *egl-13(ku194)* loss of function allele on uv1 markers. Reporter genes used are CRISPR/Cas9-engineered reporter alleles for *lin-11(ot958), unc-13(syb6325), tcd-1(syb7768)* and transgenic reporters for *flp-22(ynIs50), ocr-2(vsIs177), ocr-4(kyEx581), ida-1(vsIs269), sri-5(otEx6406)* and *NeuroPAL(otIs669)*. “ON” and “OFF” refers to reporter signal in wildtype versus *egl-13* mutant background. **B, C:** Loss and gain of utse markers in *egl-13(ku194)* mutants as seen with *exc-9(ot1866)* (panel B) and *nas-22(ot1844)* (panel C) CRISPR/Cas9-engineered reporter alleles. Animals in all panels were imaged at the young adult stage. All images are approximate lateral views with anterior to the left and ventral down. Maximum intensity projections of limited image slices are shown, which makes expression in other cells seem more variable than it really is; oftentimes the other cells are not within the image slices used. Dashed lines delineate ventral outline of worms. Scale bars 10µm. Statistical analysis was performed using Fisher’s exact test, **P < 0.01, ***P < 0.001, ****P < 0.0001.

To assess whether, like in *lin-11* mutants, the uv1 cells may adopt an utse identity in *egl-13/SoxD* mutants we used the two utse markers, *nas-22* and *exc-9*, described above. As in *lin-11* mutants, *exc-9* expression is observed in many additional cells in *egl-13* mutants (**Fig.6C**). In contrast to *lin-11* mutants, however, we observed a complete loss of *nas-22* expression in *egl-13* mutants (**Fig.6B**), indicating that *egl-13* is required not only for utse cell fusion to the anchor cell, as described before (Hanna-Rose and Han, 1999), but also for the manifestation of this specific differentiation feature of the utse cells. Taken together, since the uv1/utse precursor cells (the π cells) show additional rounds of cell division in *egl-13* mutant animals (Cinar et al., 2003), we surmise that these additional cells may adopt an utse-like fate (as inferred from *exc-9* expression), but, as inferred from the lack of *nas-22* expression, are not properly/fully differentiated utse cells. We conclude that *egl-13* is also required for uv1 differentiation and has additional effects in sister and precursor cells of the uv1 cell.

## DISCUSSION

The main purpose of our study was to probe the neuron-like features of uv1 and to characterize transcription factors that control uv1 differentiation in order to draw comparisons of uv1 to canonical neuronal differentiation programs. We observe a dichotomous gene regulatory architecture in uv1 cells that is akin to neuronal cell types: So-called “pan-neuronal” features, i.e. genes involved in synaptic vesicle release and neuropeptide processing are expressed in uv1 and their proper expression requires pan-neuronally and uv1-expressed CUT homeobox genes. In parallel, neuron/uv1-specific combinations of transcription factors control more cell-type specific aspects of neuron/uv1 differentiation. These factors appear to operate in a terminal selector-type manner, i.e. they control multiple and perhaps all cell-type specific features. The uv1 regulatory architecture involves at least one member of a family of transcription factors, homeodomain proteins, that is prominently involved in neuronal cell type specification.

One of two of the transcription factors that we characterize here as a terminal selector of uv1 differentiation through temporally controlled gene removal, *egl-38*, also has an earlier, upstream role in driving the uv1 differentiation program. *egl-38* apparently regulates the ability of the vulF cell to induce uv1 identity via an EGF-mediated signal (Chamberlin et al., 1997; Chang et al., 1999; Webb Chasser et al., 2019). Once received, EGFR signaling induces *egl-38* expression, to then drive, together with *lin-11* and *egl-13*, the terminal differentiation program of uv1. The mutant phenotypes of *lin-11* and to some extent also *egl-13* illustrate another common feature of terminal selectors that has been observed numerous times in the nervous system (Arlotta and Hobert, 2015; Reilly et al., 2022). Rather than just activating a specific differentiation program, terminal selectors often also repress alternative differentiation programs, hence resulting in cell identity changes after terminal selector removal (Arlotta and Hobert, 2015; Reilly et al., 2022). Such a cell identity change is particularly apparent in *lin-11* null mutants, in which uv1 now converts to an utse-like identity. *lin-11* may act by repressing the expression of a transcription factor that drives utse identity.

How specific combinations of terminal selectors are induced in individual neuron classes in *C. elegans* is, with a few exceptions, not well understood. uv1 fate specification requires an inductive EGF signal from the neighboring vulF cells which induces EGL-38 expression through EGF receptor (LET-23) signaling (Chang et al., 1999; Webb Chasser et al., 2019). EGL-38-independent functions of EGFR/LET-23 signaling in uv1 have been inferred (Webb Chasser et al., 2019) and those may involve the possible regulation of *lin-11* and *egl-13* expression by *let-23/EGFR*. EGF signals are required for neuroblast cell specification in several cellular contexts in *C. elegans* (Chamberlin and Sternberg, 1994; Jiang and Sternberg, 1998), but whether EGF also triggers identity specification of postmitotic neurons, as it does for the uv1 paraneuron, remains to be investigated.

One unifying theme of paraneuron development across animal phylogeny appears to be the involvement of proneural basic helix-loop-helix (bHLH) transcription factors in their initial specification (Hobert, 2025). We find the *C. elegans* ASc homolog *hlh-3* to be expressed in uv1, but its removal either in isolation or in combination with removal of the proneural Atonal homolog *lin-32* and the NeuroD1 and neurogenin homologs *cnd-1* and *ngn-1* does not affect uv1 generation. These quadruple mutant animals do, however, display partially penetrant embryonic lethality and it is possible that an absence of uv1 defects in viable “escapers” may be misleading. Another ASc-like *C. elegans* bHLH gene with proneural function, *hlh-14*, is an essential gene (Frank et al., 2003) that we could not easily include in our analysis.

Even though the involvement of bHLH factors remains unresolved, the apparent similarity in regulatory architecture of neuronal cell types and the paraneuron that we describe here suggests a potentially shared evolutionary origin of these cell types. Primitive “proto-neurons” that display a core set of signaling features –from sensory receptors to neurosecretory machinery, may have controlled such features in a modular manner. The genomic expansion of transcriptional regulators permitted the assembly of cell-type specific combinations of regulatory factors that exerted master regulatory control over the respective differentiation programs of distinct types of “proto-neurons”. Some of these cell types may have remained in a morphologically perhaps more primitive glandular type, while others evolved more elaborate morphological and functional features that resemble present-day neurons.

## ACKNOWLEDGEMENTS

We thank Chi Chen for generating *C. elegans* mutant and transgenic strains, Marion Boeglin for providing an *unc-2* reporter allele and members of the Hobert lab for comments on this manuscript. This work was funded by the NIH (R01NS039996 to OH, R24OD010943 to NES) and the Howard Hughes Medical Institute. Some strains were provided by the CGC, which is funded by NIH Office of Research Infrastructure Programs (P40 OD010440).

## DATA AVAILABILITY

Strains are available for ordering from the CGC. Digitized EM data are available at WormImage.org or upon request to NES.

